## Supplementary Information for "Active Cytoskeletal Composites Display Emergent Tunable Contractility and Restructuring"

**Contents:**

**Movie S1: Sample time-series of active actin-microtubule composites corresponding to images shown in Figure 1 of manuscript.**

**Figure S1: Plateaus in DDM image structure functions are due to myosin-driven contractile dynamics.**

**Figure S2: Active entangled actin networks exhibit faster contraction dynamics than active actin-microtubule composites.**

**Table S1: Active composites exhibit ballistic contraction.**

**Section S1: Double-Network Model and calculations**

**Figure S3: Model of an active double network with molar actin fraction $\Phi_A = 0.5$. Actin filaments are shown in magenta and microtubules are shown in green.**

**Figure S4: A single primitive cell of a kagome fiber network.**

**Section S2: Bibliography**

**Video S1. Time-series of active actin-microtubule networks show varying compositions tune dynamics and structure.** https://drive.google.com/file/d/1PlQFyRUjBFfkyA2vDHB_-k5fHO7othzW/view?usp=sharing

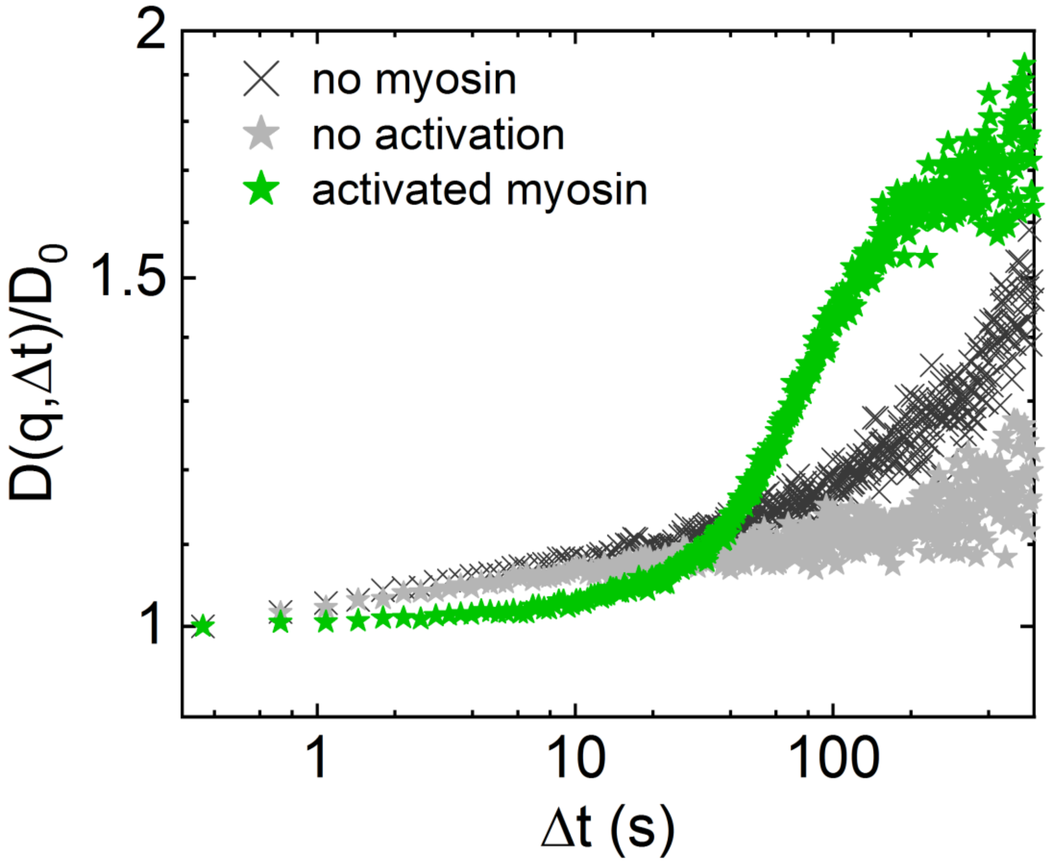


**Figure S1. Plateaus in DDM image structure functions are due to myosin-driven contractile dynamics.** Representative normalized image structure functions $\frac{D(q,\Delta t)}{D_0}$ at $q$ = 1.48 μm$^{-1}$ in the actin channel for ($\Phi_A$, $c_M$) = (0.50, 0.24) composites imaged under the following conditions: no myosin is included in the composite (dark grey crosses), myosin is included but the composite is not exposed to 488 nm light to deactivate blebbistatin (light grey stars), and myosin is included and the composite is exposed to 488nm light (green stars). Neither negative control case (light or dark grey) reaches a decorrelation plateau over the experimental time frame, demonstrating that image structure function plateaus are due to myosin-driven active dynamics.

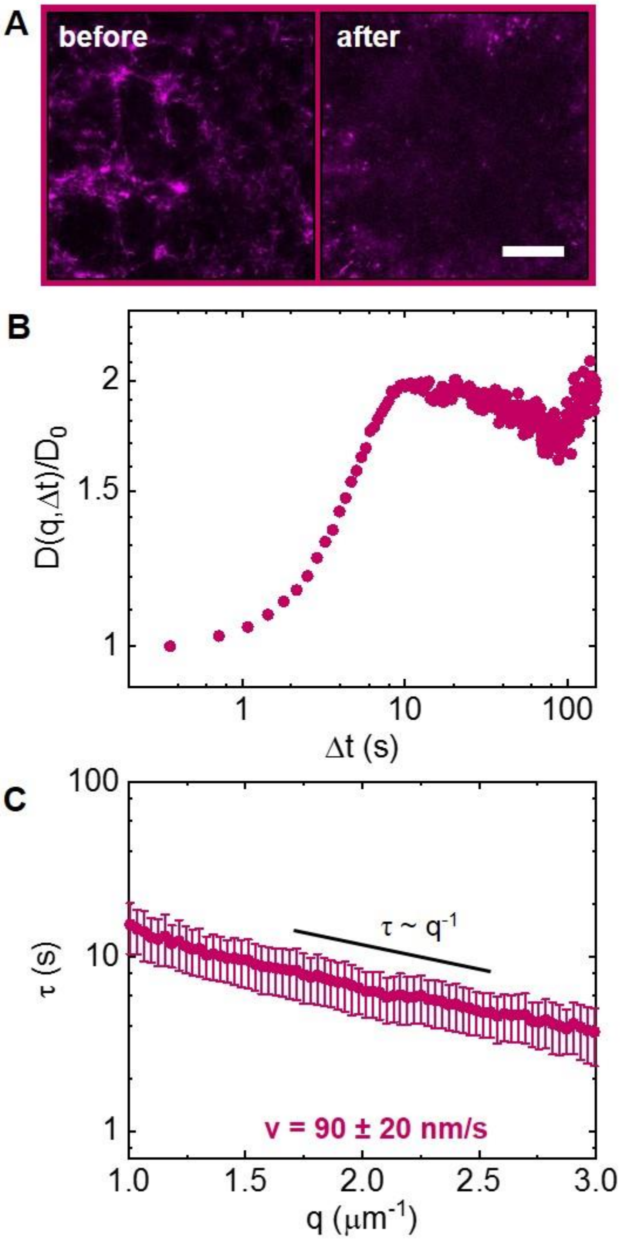


**Figure S2. Active entangled actin networks exhibit faster contraction dynamics than active actin-microtubule composites.** (A) Still images of a 5.8 μM actin network (($\Phi_A$, $c_M$)=(1, 0.24)) before and after 6 min of myosin II rearrangement. Timescale is shortened due to the network disappearing from the field of view due to faster contraction dynamics. Scale bar is 50 μm. (B) Representative normalized image structure function $\frac{D(q,\Delta t)}{D_0}$ at $q$ = 1.48 μm$^{-1}$. $\frac{D(q,\Delta t)}{D_0}$ reaches a plateau at a much earlier lag time compared to networks containing microtubules (see Fig. 2A). (C) Average $\tau(q)$ plot follows a power law relationship $\tau = \frac{1}{kq}$, from which we extract an average contraction velocity of 90 ± 20 nm/s, more than double the velocity of the ($\Phi_A$, $c_M$) = (0.75, 0.24) composite.

| ($\Phi_A$,$c_M$) | power (microtubule channel) | power (actin channel) |
|---|---|---|
| (0.25, 0.24) | -- | -- |
| (0.25, 0.48) | -0.97 ± 0.01 | -0.88 ± 0.08 |
| (0.50, 0.12) | -- | -- |
| (0.50, 0.24) | -1.00 ± 0.08 | -1.1 ± 0.1 |
| (0.50, 0.48) | -0.94 ± 0.05 | -0.87 ± 0.04 |
| (0.75, 0.12) | -1.5 ± 0.2 | -1.5 ± 0.1 |
| (0.75, 0.24) | -1.12 ± 0.07 | -1.10 ± 0.07 |

**Table S1. Active composites exhibit ballistic contraction.** For image structure functions that plateau, $\tau(q)$ plots are fit to the form $\tau \sim q^b$ over the range $q$=1-3 μm$^{-1}$, where $b$ is the power displayed in the table. Values are averaged over 3-5 time-series.

### Section S1: Double-Network Model and Calculations

#### S1a: Mathematical model for simulating myosin-driven active double-networks composed of actin filaments and microtubules

Rigidity percolation theory has been immensely successful in predicting the mechanical properties and phase transitions in single-component cytoskeletal and extracellular matrix networks as a function of filament concentrations. This theory models biopolymer networks as two interconnected networks of disordered fibers and provides a framework for connecting network rigidity to structure and composition. Here we combine rigidity percolation theory with an active double network model made of a stiff microtubule network and an active semiflexible actomyosin network.

This active rigidly percolating double-network (RPDN) is constructed as follows. Starting with two networks, each based on a fully occupied kagome lattice such that at each crosslink there are no more than two crossing fibers, we dilute the networks by uniformly and randomly removing bonds from the networks according to two different probabilities (Fig. S3). We remove bonds from the stiff microtubule network with probability $1 - p_1$ and from the semiflexible actin network with probability $1 - p_2$, where $0 < p_1, p_2 < 1$, and a contiguous series of colinear bonds constitute a fiber. The stretching moduli of the fibers in the stiff and semiflexible networks are $\alpha_1$ and $\alpha_2$ respectively, and the bending moduli are $\kappa_1$ and $\kappa_2$ respectively. The two networks interact via weak Hookean springs with spring constant $\alpha_3$, which connect the midpoints of bonds $(\boldsymbol{x_1}, \boldsymbol{x_2})$ and are only present when corresponding bonds are present in both networks. The energy cost of deforming this double network is given by:

$$E_1 = \frac{\alpha_1}{2}\sum_{<ij>} p_{1,ij}\,\left(\boldsymbol{r_{ij}} - \boldsymbol{r_{ij0}}\right)^2 + \frac{\kappa_1}{2}\sum_{<\widehat{ijk}=\pi>} p_{1,ij}\ p_{1,jk}\ \Delta\theta_{ijk}^2$$

$$E_2 = \frac{\alpha_2}{2}\sum_{<ij>} p_{2,ij}\,\left(\boldsymbol{s_{ij}} - \rho\boldsymbol{s_{ij0}}\right)^2 + \frac{\kappa_2}{2}\sum_{<\widehat{ijk}=\pi>} p_{2,ij}\ p_{2,jk}\ \Delta\beta_{ijk}^2$$

$$E_3 = \frac{\alpha_3}{2}\sum p_{1,ij}\ p_{2,ij}(\boldsymbol{x_1} - \boldsymbol{x_2})^2 \ (1)$$

where $E_1$ is the deformation energy of the stiff network, $E_2$ is the deformation energy of the semiflexible network, and $E_3$ is the deformation energy of the bonds connecting the two networks. In $E_1$ and $E_2$, the first term corresponds to the energy cost of fiber stretching, and the second term to fiber bending[1.]

In the above expression, the indices $\boldsymbol{i, j, k}$ refer to sites (nodes) in each lattice-based network, such that $p_{ij}$ is 1 when a bond between those lattice sites is present and 0 if a bond is not present. The quantities $\boldsymbol{r_{ij}}$ and $\boldsymbol{s_{ij}}$ refer to the vector lengths between lattice sites $i$ and $j$ for the deformed stiff and flexible networks respectively, while $\boldsymbol{r_{ij0}}$ and $\boldsymbol{s_{ij0}}$ are the corresponding quantities for the initial undeformed networks. Active contractility is incorporated into the semiflexible network by setting the rest length of the bonds in this network to be $\rho\boldsymbol{s_{ij0}}$, where $\rho$ is a function of myosin concentration as described later in this document and is 1 for a purely actin network and less than 1 for an actomyosin network[2]. The angles $\Delta\theta_{ijk}$ in the rigid network and $\Delta\beta_{ijk}$ in the semiflexible network correspond to the change in angles between initially collinear bond pairs $ij$ and $jk$ for the deformed and undeformed network, respectively.

Simulations of the above active RPDN model determined the linear response under 0.005% shear. We adopt a shear protocol where external deformations are applied along the top and bottom boundaries and periodic boundary conditions are used for the left and right sides of the network. For each set of parameters, active double networks containing $\sim 2 \times 10^5$ nodes were randomly generated with given fractions of bonds $1 - p_1$ and $1 - p_2$ missing. The total deformation energy was minimized for the applied macroscopic shear and the shear modulus was calculated[1] as a function of the bond occupation probabilities $p_1$ and $p_2$. The values of $p_1$ and $p_2$ were obtained from experimental molar concentrations of tubulin and actin respectively, and the contraction parameter $\rho$ was obtained using the ratio of experimental myosin and actin concentrations as described in the next section.

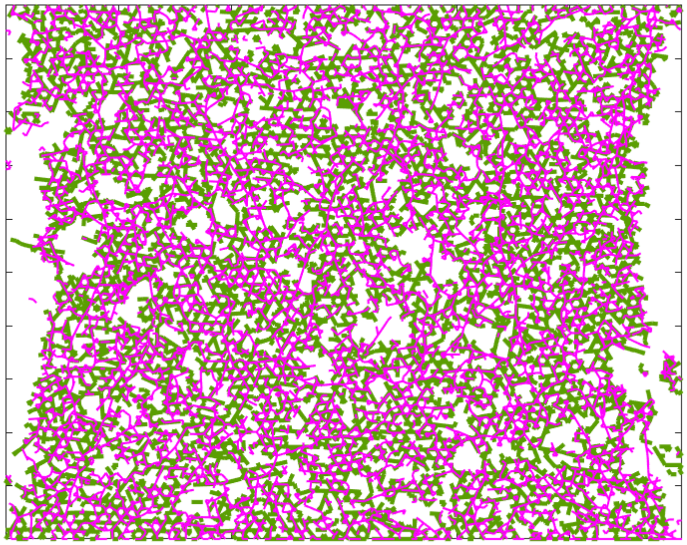

**Figure S3:** Model of an active double-network with molar actin fraction $\Phi_A$=0.5. Actin filaments are shown in magenta and microtubules are shown in green.

#### S1b: Parameter estimation informed from experiments and literature

Mesh sizes are calculated for varying composites using the equations $\eta_A = \frac{0.3}{\sqrt{c_A}}$ and $\eta_{MT} = \frac{0.89}{\sqrt{c_T}}$, which relate mesh size to polymer concentrations for actin and tubulin respectively[3]. To calculate stretching moduli and bending moduli, we use persistence lengths of 20 μm and 1 mm, and diameters of 10 nm and 50 nm, for the actin filaments and microtubules respectively[3].

By defining persistence length $l_p$ as $l_p = \frac{\kappa}{k_B T}$, where κ is the bending stiffness of the bond, $k_B$ is the Boltzmann constant and $T$ is the temperature, the bending modulus is given by the relation $\boldsymbol{\kappa} \propto \boldsymbol{l_p}$. For slender rods, the stretching modulus is given by $\boldsymbol{\alpha} \propto \frac{\boldsymbol{l_p}}{\boldsymbol{R^2}}$, where $R$ is the cross-sectional radius of the rod [4]. From these two relationships, we can calculate the stretching moduli $\alpha_1$ and

$\alpha_2$, and the bending moduli $\kappa_1$ and $\kappa_2$, of the fibers in the stiff and semiflexible networks respectively. In our simulations, all moduli are scaled by, and expressed in terms of, $\alpha_1$. To introduce contraction to the actomyosin network, we assign different values of $\rho$, or the amount the rest length of each bond is reduced due of myosin-induced contractility[2], to networks with different myosin concentrations. We use the Fermi estimate $\rho = 1 - \frac{[myosin]}{[actin]}$, such that in the absence of myosin, the network does not undergo any contraction.

To map the concentrations of actin filaments and microtubules to the bond occupation probabilities in the semiflexible and rigid networks, we use the following procedure. Let each bond in the network have length $l_0$, and let a primitive cell of the network have a length $a = l_0$, as shown in Fig. S4.

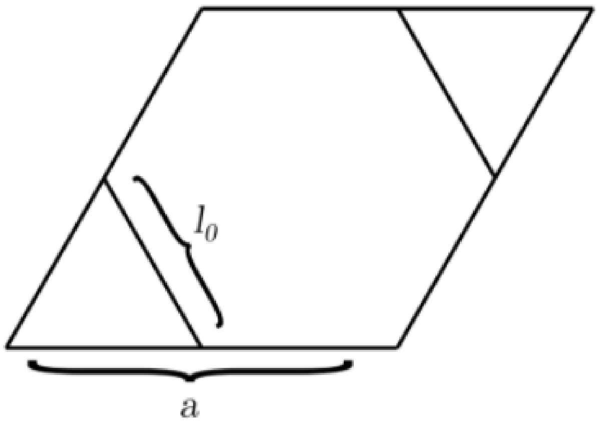


**Figure S4:** A single primitive cell of a kagome fiber network

There are eight perimeter bonds, each contributing half a bond to the primitive cell, and two interior bonds, each contributing one bond. The area of a primitive cell is $A = \frac{\sqrt{3}a^2}{2} = 2\sqrt{3}l_0^2$. For a bond occupation probability $p$, the length of fiber per unit area is then $\frac{length}{area} = \frac{6pl_0}{2\sqrt{3}l_0^2} = \frac{p\sqrt{3}}{l_0}$. Assuming the thickness of each slice of the three-dimensional sample to be $l_0$, the length per unit volume is then $\frac{length}{volume} = \frac{p\sqrt{3}}{l_0^2}$. For a given fiber type $f$, let $\lambda_f$ be the number of monomers per unit length. If the molar concentration is $[f]$, then the total fiber length per unit volume should be

$\frac{length}{volume} = \frac{[f]N_A}{\lambda_f}$ where $N_A$ is Avagadro's constant $\approx 6.022 \times 10^{23}$. By setting the two length per volume equations equal to each other, $\frac{[f]N_A}{\lambda_f} = \frac{p\sqrt{3}}{l_0^2}$, we find $p = \frac{l_0^2[f]N_A}{\lambda_f\sqrt{3}}$.

To determine $\lambda_f(actin)$ and $\lambda_f(microtubule)$, which represent the number of monomers per unit length for actin filaments and microtubules, respectively, we use 2.7 nm per monomer for actin filaments[5] and 12 nm per 13 tubulin dimers for microtubules[6]. In using the above-mentioned calculation to map the microtubule and actin fractions to the bond occupation probabilities $p_1$ and $p_2$ respectively, we have set each bond occupation probability to 1 when the corresponding concentration is 5.8 μM.

#### S1c: Creating strain maps of actin and microtubule networks

To produce strain maps, we begin by producing a Delaunay triangulation[7] of the vertices of the network in its undeformed state and compute the area of each triangular facet. After the bonds of the actin network are subjected to contractile forces, and the network is relaxed to its energetic ground state, we use the new positions of the vertices in the microtubule and actin networks to produce deformed triangular meshes. We compute the new area of each deformed triangular mesh cell, subtract from this area the area of the corresponding, undeformed mesh cell, and divide by the area of the undeformed cell to find relative change in area. We next identify all groups of contiguous cells that undergo contraction, and all contiguous groups that undergo extension. To color-code contiguous regions that undergo contractile or extensile deformation, we take the fourth root of each relative extension and contraction to account for the large dynamic range in contraction and extension. We map these adjusted values to a color gradient in which deep red regions are highly contractile, and deep blue values are highly extensile.
